## Supplemental files for "Expression of varicella-zoster virus VLT-ORF63 fusion transcript induces broad viral gene expression during reactivation from neuronal latency"

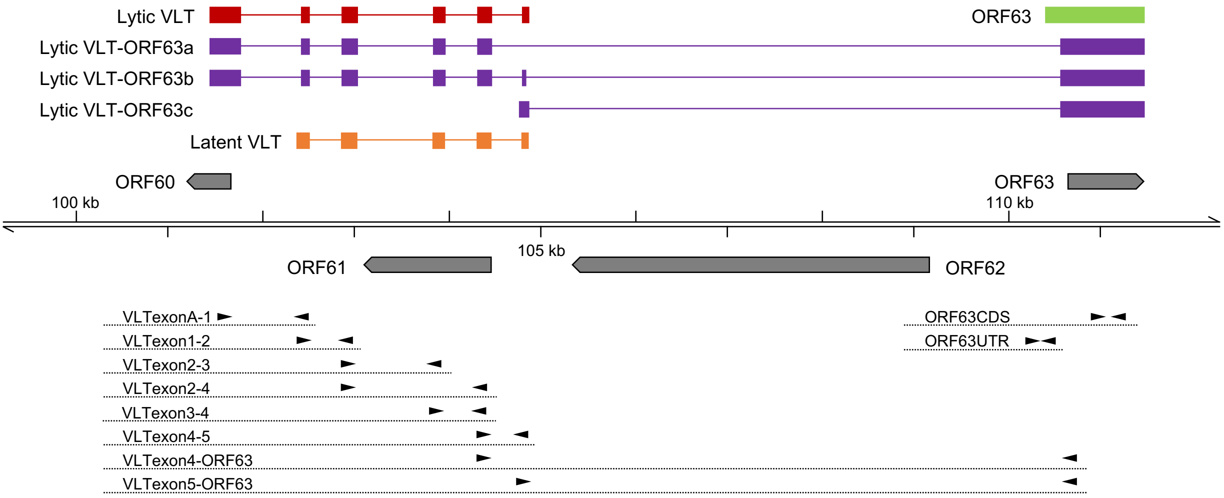


***Supplementary Figure 1. Location of primer sets detecting transcripts from VLT and ORF63 loci.*** Schematics of major transcripts from VLT and ORF63 loci are shown in color: lytic VLT isoform (red), latent VLT isoform (orange), canonical ORF63 (light green) and lytic VLT-ORF63 isoforms (purple). Location of primer sets used for RT-qPCR analysis to detect transcripts from VLT and ORF63 loci are depicted.

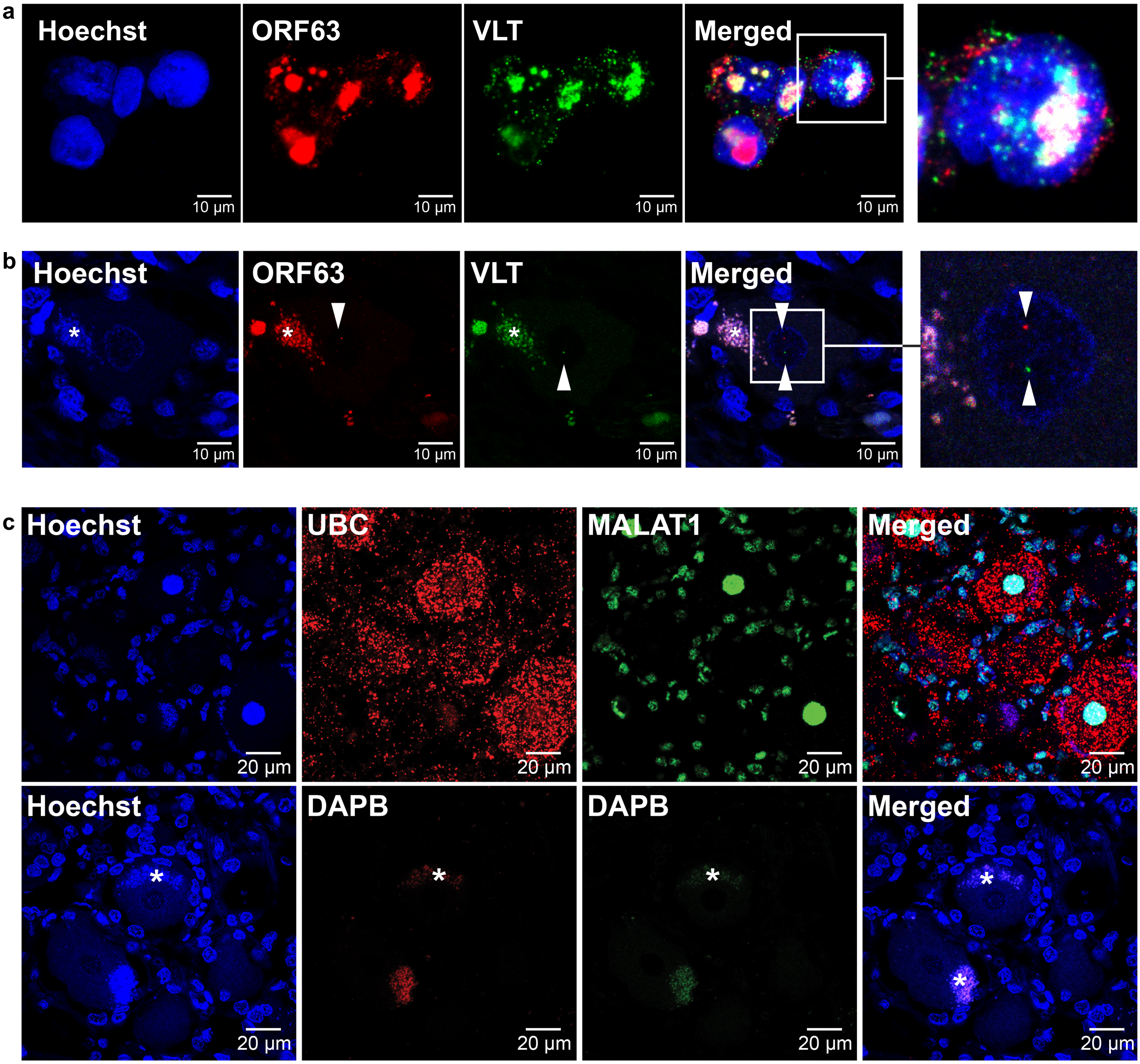


***Supplementary Figure 2. Detection of VZV ORF63 and VLT RNA by fluorescent multiplex in situ hybridization.*** Detection of VZV ORF63 and VLT RNA by multiplex fluorescent *in situ* hybridization (mFISH) on **a** lytically VZV-infected ARPE-19 cells and **b** latently VZV-infected human TG. **a, b** Right panels represent enlargements of area indicated by white box. **b** Rare detection of ORF63 (red) and VLT (green) as discrete puncta in nuclei of human TG neurons. Asterisks indicate autofluorescent lipofuscin granules in neurons and arrowheads indicate ORF63 and/or VLT mFISH signal. **c** Human TG were stained with mFISH using positive [UBC (red) and MALAT1 (green)] and negative control probes (DAPB, green) to demonstrate specificity of the mFISH assay and RNA integrity of the tissue assayed. Nuclei were stained with Hoechst (blue).

**pVLT**

MPRLLRDRIA GIPNRVRTYQ GAVFTPWVPD IPTLTTNSNT QILDDHGSPA PRSGVAVQIQ SSHTPPGSPI EQQDGLHWTP AERTLDAGGG PCPNTNKAEV VQTRHGFSEI GNGAHAYGAD KERYEDISPP PCNTRK

**pORF63**

MFCTSPATRG DSSESKPGAS VDVNGKMEYG SAPGPLNGRD TSRGPGAFCT PGWEIHPARL VEDINRVFLC IAQSSGRVTR DSRRLRRICL DFYLMGRTRQ RPTLACWEEL LQLQPTQTQC LRATLMEVSH RPPRGEDGFI EAPNVPLHRS ALECDVSDDG GEDDSDDDGS TPSDVIEFRD SDAESSDGED FIVEEESEES TDSCEPDGVP GDCYRDGDGC NTPSPKRPQR AIERYAGAET AEYTAAKALT ALGEGGVDWK RRRHEAPRRH DIPPPHGV

**pVLT-ORF63**

MPRLLRDRIA GIPNRVRTYQ GAVFTPWVPD IPTLTTNSNT QILDDHGSPA PRSGVAVQIQ SSHTPPGSPI EQQDGLHWTP AERTLDAGGG PCPNTNKAEV VQTRHGFSEI GNGAHAYGAG FVRFITRQRR VGFKGKGYYG PKDMFCTSPA TRGDSSESKP GASVDVNGKM EYGSAPGPLN GRDTSRGPGA FCTPGWEIHP ARLVEDINRV FLCIAQSSGR VTRDSRRLRR ICLDFYLMGR TRQRPTLACW EELLQLQPTQ TQCLRATLME VSHRPPRGED GFIEAPNVPL HRSALECDVS DDGGEDDSDD DGSTPSDVIE FRDSDAESSD GEDFIVEEES EESTDSCEPD GVPGDCYRDG DGCNTPSPKR PQRAIERYAG AETAEYTAAK ALTALGEGGV DWKRRRHEAP RRHDIPPPHG V

**pORF63-N+**

MDCTGHRQRG HWTLVEVHAR IQTKQKSCKH GMVFPRSETV LMHMVQIKSD TKTFLHPPVI PVNKGFVRFI TRQRRVGFKG KGYYGPKDMF CTSPATRGDS SESKPGASVD VNGKMEYGSA PGPLNGRDTS RGPGAFCTPG WEIHPARLVE DINRVFLCIA QSSGRVTRDS RRLRRICLDF YLMGRTRQRP TLACWEELLQ LQPTQTQCLR ATLMEVSHRP PRGEDGFIEA PNVPLHRSAL ECDVSDDGGE DDSDDDGSTP SDVIEFRDSD AESSDGEDFI VEEESEESTD SCEPDGVPGD CYRDGDGCNT PSPKRPQRAI ERYAGAETAE YTAAKALTAL GEGGVDWKRR RHEAPRRHDI PPPHGV

***Supplementary Figure 3. In silico translation of VLT, canonical ORF63 and VLT-ORF63 fusion transcripts.*** Amino acid (aa) sequence of VLT protein (pVLT; encoded by VLT or VLT-ORF63b), ORF63 protein (pORF63; encoded by ORF63 or VLT-ORF63c), VLT-ORF63 protein (pVLT-ORF63; encoded by VLT-ORF63a) and pORF63-N+ (encoded by VLT-ORF63b). The sequence of 24 aa peptide used as immunogen to generate the chicken anti-pVLT-ORF63 polyclonal antibody is highlighted in grey color.

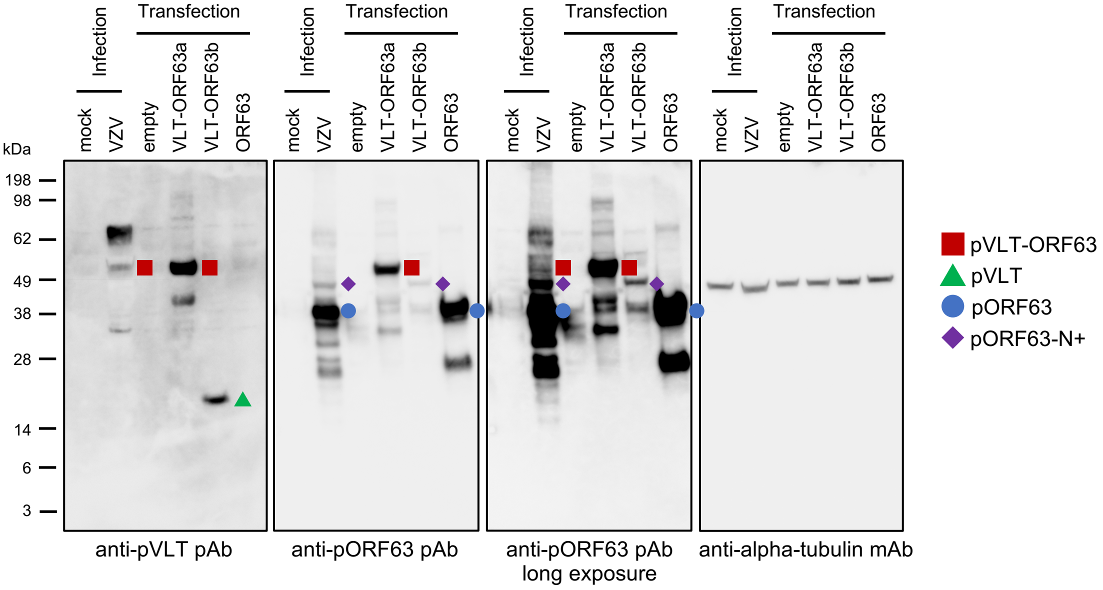


***Supplementary Figure 4. Protein coding potential of VLT-ORF63 fusion transcripts.*** Immunoblotting analysis using antibodies directed to pVLT and pORF63 in the context of mock- or VZV-infection in CS-CA-empty or CS-CA-VZV transfected ARPE-19 cells. Red squares indicate pVLT-ORF63 (45.989-kDa), green triangles indicate pVLT (14.728-kDa), blue circle indicates pORF63 (30.494-kDa) and purple diamonds indicate pORF63-N+ (40.723-kDa). Images are representative of two independent experiments. Molecular weight marker (kDa) is shown in left.

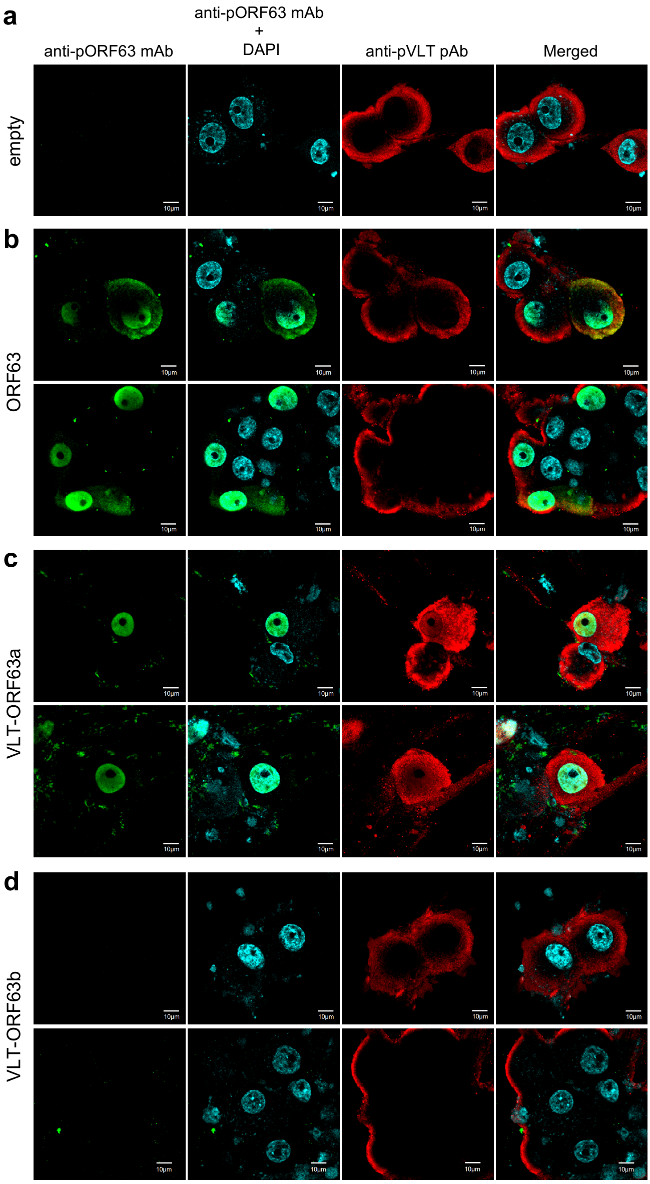


***Supplementary Figure 5. Transduced HSN express ectopic proteins encoded by replication incompetent lentivirus vectors.*** Confocal analysis using antibodies against pORF63 and pVLT in HSN transduced with replication incompetent lentivirus vectors encoding the following ORFs: **a** no gene (empty), **b** ORF63, **c** VLT-ORF63a or **d** VLT-ORF63b. HSN cultures were maturated for 49 days and transduced with each vector for 14 days. The signal obtained with anti-pORF63 mAb staining was observed in the nucleus of both ORF63- and VLT-ORF63-transduced HSN, in cytoplasm of ORF63-transduced HSN, but undetectable in HSN transduced with empty vector or VLT-ORF63b. The anti-pVLT pAb showed a nonspecific cytoplasmic signal in all transduced HSN, including 'empty', and was too strong to determine if pVLT alone is expressed by VLT-ORF63b transduction. Nuclear specific signal by anti-pVLT pAb was only detected in VLT-ORF63a-transduced HSN and co-localized with the pORF63 signal, indicating nuclear expression of pVLT-ORF63.

**Supplementary Table 1. Primers for qPCR assay.**

| Target | Name | Sequence (5' → 3') |
|---|---|---|
| beta-actin | beta-actinF961 | GCA CCC AGC ACA ATG AAG A |
| | beta-actinR1024 | CGA TCC ACA CGG AGT ACT TG |
| VLT, VLT-ORF63 | VLTexonA101426F | CAA CGG AGT GTC GTC TTG GA |
| | VLTexon1F102413 | GGC ATT TTA AAC GGG TCC GG |
| | VLTexon2F102837 | CGA GAC CGG ATT GCG GGC AT |
| | VLTexon3F103794 | TGG ACG ATC ACG GTA GTC CT |
| | VLTexon4F104342 | AAC ACG GCA TGG TTT TTC CG |
| | VLTexon5F104763 | ACG AAG ACA TTT CTC CAC CCC |
| | VLTexon1R102394 | CCG GAC CCG TTT AAA ATG CC |
| | VLTexon2R102864 | CCC TGG TAA GTC CGT ACA CG |
| | VLTexon3R103847 | ATT GAA TCT GCA CAG CAA CCC |
| | VLTexon4R104361 | CGG AAA AAC CAT GCC GTG TT |
| | VLTexon5R104778 | ACC CTC GAG TAC GGG TAT TAC AGG G |
| | ORF63R19 | CCG GTG AGG TGC AAA ACA TG |
| ORF4 | ORF4F363 | GGG GAC ATC GAC GAT CAT CC |
| | ORF4R415 | GTA GGA CGC CGT CTT CGA TT |
| ORF9 | ORF9F224 | AAA AAT ACG ACC CCT CGC GT |
| | ORF9R298 | TCA TGT CTC AAA CGG GCC TC |
| ORF16 | ORF16F1091 | GGA AAC TCC CCG AAA CCA |
| | ORF16R1158 | CAC TGG AGG AGC CAC ACA A |
| ORF29 | ORF29F2381 | GCC TTG CAA GTG CGT ACC |
| | ORF29R2440 | CTA GGG CCC CGT GTA ACA TA |
| ORF31 | ORF31F331 | CAG GAC GCC GAA ACA AAA |
| | ORF31R393 | TAC GAT TGT GGA GCC TGT TG |
| ORF49 | ORF49F21 | CGG TCG AGG AGG AAT CTG TG |
| | ORF49R80 | CCG TTG CAC GTA ACA AGC TC |
| ORF61 | ORF61F150 | CAG CGT CCA GTG TCC TCT CT |
| | ORF61R210 | ACT TAC GAT CTT ATG CAG GAT GG |
| ORF62 | ORF62F2016 | TCC ACC GGA TGA TCG TTT AC |
| | ORF62R2083 | GGA GGC TTC TGC TCT CGA C |
| ORF63 CDS | ORF63F556 | TCG GAC GGG GAA GAC TTT AT |
| | ORF63R622 | CGT CTG GTT CAC AAG AAT CG |
| ORF63 5'-UTR | ORF63up100F | AAC GTT TGG GTG TGT GTT TTG T |
| | ORF63up1R | GTC CTT GGG GCC GTA GTA AC |
| ORF66 | ORF66F688 | TTC CCC GTG GAT ATT AAT GC |
| | ORF66R752 | GGA GAG TTT GTG GCG ATT GT |
| ORF68 | ORF68F661 | TTA AAA CAT ACA ACA TGC TTT CAA GA |
| | ORF68R720 | AGT ATT TTC CGC GCA ATC C |
| CS-CA vector | CSCA1831F | CAA CTC ACA GTC TGG GGC AT |
| | CSCA1969R | TAG CAT TCC AAG GCA CAG CA |

**Supplementary Table 2. Clinical features of human trigeminal ganglia donors used for RNA extraction.**

| Donor[a] | Age | Gender[b] | Cause of death | Neurological Disease | PMI[c] (hr:min) |
|---|---|---|---|---|---|
| 1 : s07/122 | 94 | F | Cerebrovascular accident | Non-demented control | 4:05 |
| 2 : s09/066 | 99 | F | unknown | Non-demented control | 4:15 |
| 3 : s09/185 | 70 | M | Cachexia and dehydration | Alzheimer's disease | 4:00 |
| 4 : s10/349 | 95 | F | Cachexia | Alzheimer's disease | 4:30 |

[a]Donor number in this study and corresponding Netherlands Brain Bank number.
[b] F, female; M, male. [c]PMI: post-mortem interval.

**Supplementary Table 3. Clinical features of human trigeminal ganglia donors used for ISH.**

| Donor[a] | Age | Gender[b] | Cause of death | Neurological Disease | PMI[c] (hr:min) |
|---|---|---|---|---|---|
| s06/235 | 80 | F | Cardiac arrest | Bipolar disorder | 9:30 |
| s11/084 | 63 | F | Gastrointestinal bleeding | Frontotemporal dementia; Pick's disease | 4:00 |
| s11/088 | 81 | F | Cachexia and dehydration | Alzheimer's disease | 3:35 |
| s11/092 | 64 | F | Cachexia | Frontotemporal dementia; tauopathy | 7:30 |
| s11/097 | 90 | M | Cardiac insufficiency | Lewy body dementia | 4:05 |
| s13/010 | 89 | F | Heart failure with dehydration | Non-demented control | 6:35 |
| s13/013 | 89 | M | Urosepsis | Non-demented control | 6:50 |

[a]Donor number in this study and corresponding Netherlands Brain Bank number.
[b] F, female; M, male. [c]PMI: post-mortem interval.

**Supplementary Table 4. Primers for cDNA cloning and 5'-RACE analysis.**

| Target | Name | Sequence (5' → 3') |
|---|---|---|
| **cDNA cloning** | | |
| CS-CA-MCS linearization | CSCAInFusionF | AGA TAT CCA GCA CAG TGG CG |
| | CSCAInFusionR | CGT TGCCCA GGA GCT GTA GG |
| CS-CA-ORF63 | ORF63up20xhoF | ACC CTC GAG GTT ACT ACG GCC CCA AGG |
| | ORF63xhoR | ACC CTC GAG CTA CAC GCC ATG G |
| CS-CA-VLT-ORF63 | InFusionCSCAVLTcoreTSSF | CCT ACA GCT CCT GGG CAA CGG CAG ACT ATC CAG TTG GCA |
| | InFusionCSCAORF63R | CGC CAC TGT GCT GGA TAT CTC TAC ACG CCA TGG |
| **5'-RACE analysis** | | |
| First-strand cDNA synthesis | SMARTerIIA Oligonucleotide | AAG CAG TGG TAT CAA CGC AGA GTA CAT GGG XXXXX (X :=undisclosed base) |
| | 5'-RACE CDS Primer A | $(T)_{25}$VN (V=A, G or C, N=A, C, G or T) |
| 5'-RACE PCR | Universal Primer Long | CTA ATA CGA CTC ACT ATA GGG CAA GCA GTG GTA TCA ACG CAG AGT |
| | Universal Primer Short | CTA ATA CGA CTC ACT ATA GGG C |
| | (InFusion)VLTexon4R104361 | GAT TAC GCC AAG CTT CGG AAA AAC CAT GCC GTG TT |
| | (InFusion)VLTexon5R104799 | GAT TAC GCC AAG CTT GTT TGT GGA CTT ACC TTT ATT TAC G |
| | (InFusion)ORF63R622 | GAT TAC GCC AAG CTT CGT CTG GTT CAC AAG AAT CG |
| | (InFusion)ORF63R805 | GAT TAC GCC AAG CTT GGC GCG GGG CTT CGT GTC GA |
| pRACE linearization | pRACE-F | AAG CTT GGC GTA ATC ATG GTC |
| | pRACE-R | AGT GAG TCG TAT TAG GAA TTC AC |
| Sequencing | M13forward | GCC GCT GTA AAA CGA CGG CCA GT |
| | M13reverse | GGC CGC AGG AAA CAG CTA TGA CC |
